# Involvement of orexin nerves in early stage of Alzheimer’s disease model mice and preventive effect of orexin receptor antagonists

**DOI:** 10.1101/2025.01.28.635390

**Authors:** Kazuhiro Hada, Yuki Murata, Ohi Yoshiaki, Sana Hashimoto, Hinata Watanabe, Kayoko Ozeki, Takaomi C Saido, Takashi Saito, Hiroki Sasaguri, Hiroyuki Mizoguchi, Kiyofumi Yamada, Yoshifumi Wakiya

**Affiliations:** Laboratory of Pharmacy Practice and Sciences, School of Pharmacy, Aichi Gakuin University; Department of Neuropsychopharmacology and Hospital Pharmacy, Nagoya University Graduate School of Medicine; Laboratory of Organic and Medicinal Chemistry, School of Pharmacy, Aichi Gakuin University; Laboratory of Neuropharmacology, School of Pharmacy, Aichi Gakuin University; Laboratory for Proteolytic Neuroscience, RIKEN Center for Brain Science; Department of Neurocognitive Science, Institute of Brain Science, Nagoya City University Graduate School of Medical Sciences; Department of Neuroscience and Pathobiology, Research Institute of Environmental Medicine, Nagoya University; Dementia Pathophysiology Collaboration Unit, RIKEN Center for Brain Science; Division of Behavioral Neuropharmacology, International Center for Brain Science (ICBS), Fujita Health University

**Keywords:** Aβ, Alzheimer’s disease, cognition, mild cognitive impairment, orexin, orexin receptor antagonist, lemborexant, suvorexant

## Abstract

**Background:** Mild cognitive impairment (MCI) is a condition between healthy cognition and dementia with a high probability of progression to Alzheimer’s disease (AD). Elevated levels of orexin (OX) have been reported in the cerebrospinal fluid of patients with MCI and AD.

**Objective:** To investigate the efficacy of dual OX receptor antagonists (suvorexant and lemborexant) in an early-stage AD mouse model (App-KI).

**Methods:** The expression of the *OX receptor* gene, the levels of suvorexant and lemborexant in the brain after a single oral dose, and their effects on locomotor activity were investigated in wild-type mice. In addition, the cognitive function of wild-type and App-KI mice was assessed using the Y-maze test after oral administration of suvorexant or lemborexant once a day for 60 d. After the behavioral test, amyloid-beta levels were quantified in the hippocampal CA1 region of App-KI mice.

**Results:** The expression of the *OX receptor* gene was highest in the lateral hypothalamus and was also observed in other brain regions. Drug levels peaked at 20–40 min for suvorexant and 15 min for lemborexant and declined, still detectable 24 h later. Locomotor activity was reduced after suvorexant or lemborexant administration; however, 24 h after administration, locomotor activity did not differ from the control group. In particular, the Y-maze test showed that suvorexant and lemborexant prevented cognitive impairment in App-KI mice. Furthermore, suvorexant and lemborexant suppressed amyloid-beta accumulation in the hippocampal CA1 region of App-KI mice.

**Conclusion:** The results suggest that suvorexant and lemborexant effectively suppress the early stages of AD.

## Introduction

Mild cognitive impairment (MCI) is an intermediate state between healthy cognition and dementia and is considered a predementia condition that is likely to progress to Alzheimer’s disease (AD).^1^ The main symptom of MCI is memory loss. However, this condition has little impact on daily life and cannot be diagnosed as dementia.^2^ Amyloid-beta (Aβ) and total- and phosphorylated-tau protein levels are elevated in the cerebrospinal fluid and its accumulation in the brain parenchyma; however, the senile plaques and abnormal phosphorylation of the tau protein are mild.^3^ Acetylcholinesterase inhibitors, NMDA-type glutamate receptor antagonists, and anti-amyloid monoclonal antibodies are used to treat AD. Acetylcholinesterase inhibitors and NMDA-type glutamate receptor antagonists are symptomatic treatments that may slow disease progression, but only slightly prolong prognosis and are not fully effective.^4^ Recanemab, an anti-amyloid monoclonal antibody, is effective in removing amyloid but should be used with caution, as deaths due to amyloid-related imaging abnormalities have been reported as a side effect.^5–7^ The appropriate prevention and treatment of MCI may prevent or delay AD onset. However, the lack of effective therapeutic agents and the absence of reliable treatment methods require a new approach from a new perspective.

In recent years, several clinical studies have reported elevated levels of orexin (OX) in the cerebrospinal fluid of patients with MCI and AD,^8,9^ suggesting an association between OX, MCI, and AD. OX is a neuropeptide that was discovered in 1998. Two subtypes of OXs have been reported, OXA and OXB, which are enzymatically generated from prepro-orexin. OXA has a higher affinity for OX 1 receptor (OX1R) than OX 2 receptor (OX2R), while OXB has a similar affinity for OX1R and OX2R.^10^ OX neuron cell bodies are located in the lateral hypothalamic area (LHA) and project to several brain regions, including the cortex and hippocampus (Hip).^11,12^ Although much is still unknown due to the short time since its discovery, it contributes to feeding behavior and the maintenance of wakefulness and sleep regulation in the brain, and its relationship to psychiatric and neurodegenerative disorders such as narcolepsy, depression, ischemic stroke, and AD, which may coexist with sleep disturbance, has attracted attention.^8–10,13–25^ Among these, the activation of the OX system has been reported to be positively correlated with the accumulation of Aβ and tau, leading to sleep disorder in animal models of AD.^26^ This makes them potential therapeutic targets for AD.^27^

The OX receptor antagonists suvorexant and lemborexant have been approved as hypnotic medications in the USA and Japan, respectively. Suvorexant has a similar affinity for OX1R and OX2R, while lemborexant has almost 15-fold higher affinity for OX2R than OX1R.^28,29^ Recently, suvorexant and almorexant, dual OXR antagonists, were reported to inhibit cognitive impairment and brain Aβ accumulation in patients with AD and mouse models.^30–33^ However, administration of OX1R-selective antagonists to AD model mice has been found to exacerbate cognitive impairment.^34,35^ Meanwhile, OX2R-selective antagonists have been reported to suppress cognitive impairment and AD pathology in mouse models of AD.^36^ Therefore, antagonizing OX2R may be effective in ameliorating AD pathology and cognitive impairment; however, no studies have reported on lemborexant, which has an almost 15-fold higher affinity for OX2R.

In recent years, the relationship among OX, MCI, and AD has become clear; however, more evidence is needed to confirm its clinical application. Furthermore, the involvement of OX neurons in early-stage AD mouse models remains controversial. Therefore, we investigated (i) *OX1R* and *OX2R* gene expression in the target brain regions; (ii) the pharmacokinetics of suvorexant and lemborexant in the Hip, orbitofrontal cortex (OFC), and LHA; and (iii) their effects on locomotor activity in wild-type (WT) mice. In addition to (iv) the effect of repeated administration of suvorexant or lemborexant on cognitive impairment and (v) on Aβ accumulation in the early-stage AD mouse model to determine whether suvorexant and lemborexant are effective in the early stages of AD.

## Materials and Methods

### Animals

*App^NL-G-F/NL-G-F^* (C57BL/6-App<tm3(NL-G-F)Tcs>; App-KI) mice were generated in a C57BL/6J genetic background and provided by the RIKEN BRC through the National BioResource Project of the MEXT, Japan.^37,38^ App-KI mice and their WT littermates were obtained by breeding heterozygous *App^+/NL-G-F^* mice. Male and female mice aged 4−6 months were used in this experiment (male WT = 147, male App-KI = 56, female WT = 45, and female App-KI = 44). Mice were housed at a density of 5−6 mice/cage (28 cm × 17 cm × 13 cm) under standard conditions (23 ± 1°C, 50 ± 5% humidity) with a 12 h light/dark cycle. Food and water were provided ad libitum. All animal experiments were approved and performed according to the Animal Care and Use Regulations of the School of Pharmacy, Aichi Gakuin University, Nagoya, Japan.

### Gene expression analysis

Gene expression analysis was performed as previously described with minor modifications.^39,40^ Four 6-month-old male WT mice were sacrificed for gene expression analyses. Hip, OFC, cerebral cortex, striatum, nucleus accumbens, and LHA tissues were rapidly dissected on ice and total RNA was extracted using the RNeasy Mini Kit (Qiagen, Hilden, Germany). Total RNA was reverse transcribed to cDNA using ReverTra Ace^®^ qPCR RT Master Mix (TOYOBO, Osaka, Japan). The expression of the *OX1R* and *OX2R* genes was measured from 500 ng of cDNA using the THUNDERBIRD^®^ SYBR^TM^ qPCR Mix (TOYOBO) and the StepOnePlus^TM^ Real-Time PCR System Upgrade (Thermo Fisher Scientific, MA, USA). The *Gapdh* gene was used as a housekeeping gene. In this study, the following primers were used: *mOX1R* forward: TTGGTGCGGAACTGGAAAC; *mOX1R* reverse: CCATCAGCATCTTAGCCGTCT; *mOX2R* forward: TTCCCGGAACTTCTTCTGTGG; *mOX2R* reverse: TCAGCAGCAACAGCGCTAATC; *Gapdh* forward: CAATGTGTCCGTCGTGGATCT; *Gapdh* reverse: GTCCTC AGTGTAGCCCAAGATG.

### Drug preparation

Belsomra^®^ (MSD, NJ, USA) or Debigo^®^ (Eisai, Tokyo, Japan) tablets were suspended in 400 mL of water and stirred at 65°C for 10 min; 400 mL of ethyl acetic acid or 400 mL of chloroform was added to the Belsomra^®^ or Debigo^®^ suspension, respectively. The solution was filtered through a celite pad and the filtrate was extracted with 200 mL of ethyl acetate for Belsomra^®^ or 200 mL of chloroform for Debigo^®^, respectively, three times. The organic phase was then dried over MgSO_4_ and concentrated under pressure. The residue was purified using an Isolera^TM^ One (Biotage, Uppsala, Sweden) to obtain suvorexant or lemborexant.

### LC-MS/MS analysis

Qualitative and quantitative determination of suvorexant and lemborexant by LC-MS/MS was performed as previously described with slight modifications.^41–44^ Suvorexant and lemborexant were determined in positive ion mode using an LC system (Nexera 2; Shimadzu, Kyoto, Japan) connected to a triple quadrupole mass spectrometer (LC-MS 8040; Shimadzu) with an electrospray ion source. LC separation of suvorexant, lemborexant, and estazolam-*d5*, internal control, was performed using a C18 column (2.1 × 50 mm, 3 μm particle size; GL Sciences, Tokyo, Japan) maintained at 35°C. Separation was performed by gradient elution with mobile phases A (0.1% formic acid in water; MPA) and B (0.1% formic acid in acetonitrile; MPB). The gradient program was run as follows: 40% MPB (linear gradient, v/v) at 0 min, 40−90% MPB at 4 min, 90% MPB at 2 min, and 90−40% MPB at 2 min at a flow rate of 0.2 mL/min with a run time of 8 min for each 3 μL injection volume. Multiple reaction monitoring mode was applied for the detection of suvorexant, lemborexant, and estazolam-*d5* by monitoring the transitions: *m/z* 451.4 to 186.1 for suvorexant, *m/z* 411.4 to 287.1 for lemborexant, and *m/z* 300.25 to 272.1 for estazolam-*d5*. Details of the LC-MS/MS conditions used for the measurements are provided in Table 1.

**Table 1.**
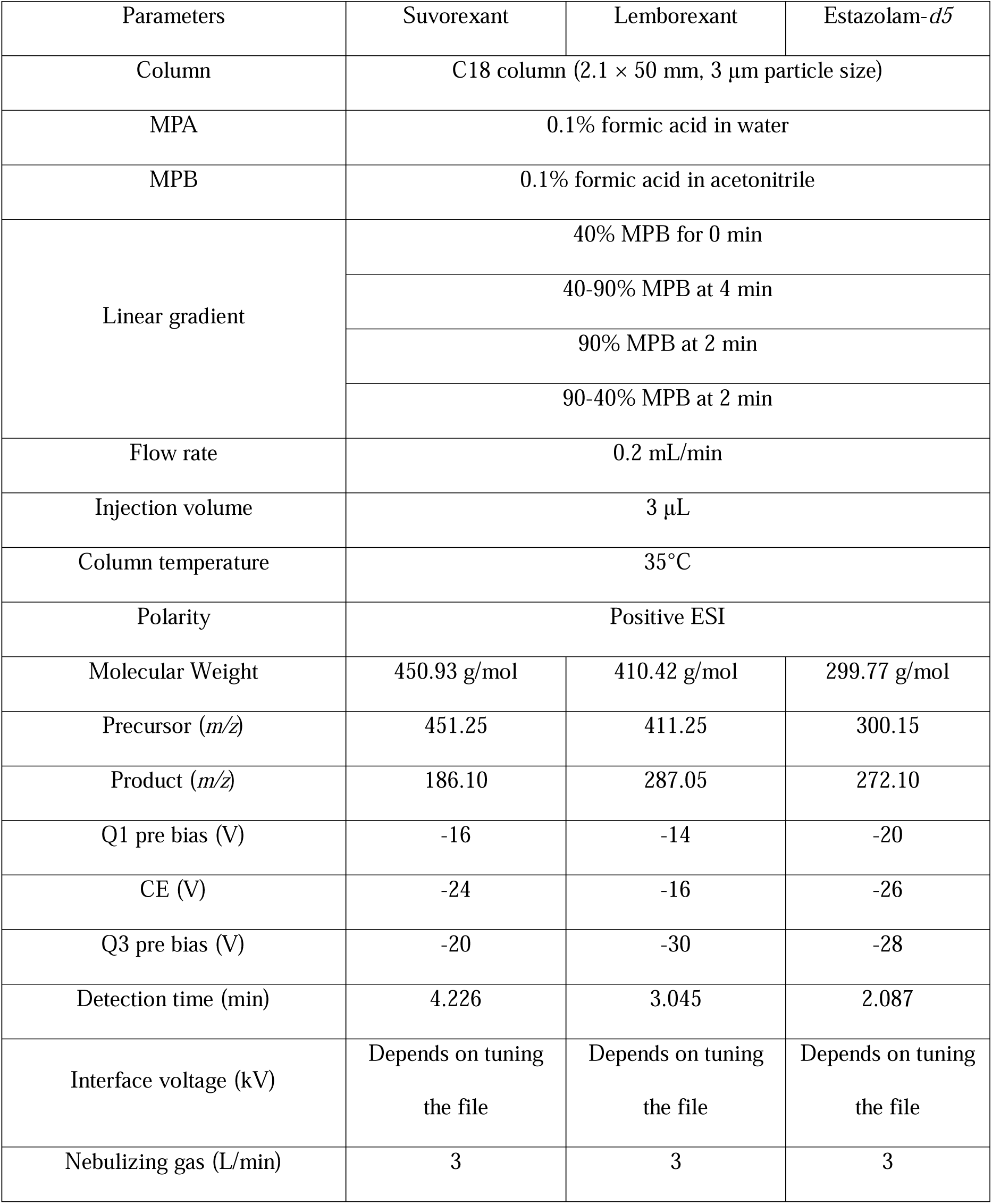
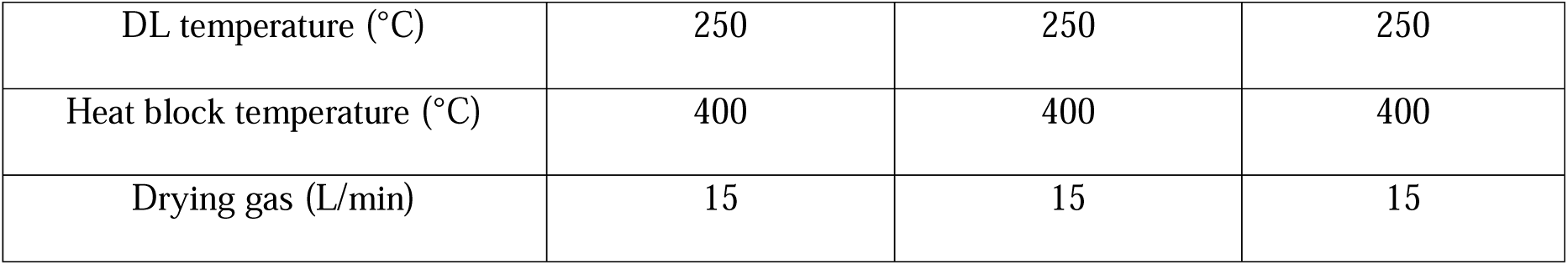
Summary of optimized parameters for LC-MS/MS analysis.

### Drug treatment

Purified suvorexant and lemborexant were assayed for purity using LC-MS/MS, and only those with >95% purity were used in the experiments. Suvorexant and lemborexant were suspended in 1% sodium carboxymethylcellulose powder (Sigma-Aldrich) dissolved in saline at 3.0 mg/mL. Vehicle, suvorexant, and lemborexant were administered orally (10 mL/kg) once or repeatedly once daily using an oral sonde (Fuchigami, Kyoto, Japan).

### Measurement of drug levels in the brain using an LC-MS/MS

Seventy 6-month-old male WT mice received a single oral dose of suvorexant (30 mg/kg) or lemborexant (30 mg/kg). Five mice were sacrificed after 10, 20, 30, 40, and 60 min, 12 h, and 24 h after receiving suvorexant and after 5, 10, 15, 30, and 60 min, and 12 h, and 24 h after receiving lemborexant. Hip, OFC, and LHA tissues were rapidly dissected on ice according to the Mouse Brain Atlas (Franklin and Paxinos, 1997) using 2 mm diameter biopsy punches (Integra Miltex, NH, USA). The tissue was then sonicated in a 120 μL MPA + MPB solution (MPA: MPB = 1:1). After sonication, the samples were added to 100 μL of MPB to remove the protein and centrifuged at 17,000 g for 20 min. The supernatant was collected and stored at - 80°C until use. The samples were spiked with estazolam-*d5* (final concentration 500 ng/mL) at the time of measurement. Brain suvorexant and lemborexant levels were measured in triplicate using LC-MS/MS under the same conditions as those used to confirm the purity of suvorexant and lemborexant. The areas under the curve (AUC)_10-60_ and AUC_5-60_ were calculated using the linear trapezoidal rule with extrapolation.

### Locomotor activity

The locomotor test was performed as described in a previous report with minor modifications.^45^ Twenty-seven 6-month-old male WT mice were randomly divided into three groups (WT vehicle, WT suvorexant, WT lemborexant). Each mouse was placed in a standard transparent rectangular rodent cage under normal light conditions (380−400 lx). The measurements were taken immediately and 24 h after administration for 120 min in each mouse. Locomotor activity was measured using an infrared sensor (WO-03; Brain Science Idea, Osaka, Japan) and a receiver box (DIG-807; Med Associates Inc., VT, USA). Data were analyzed using data acquisition

### Y-maze

The Y-maze test was performed as previously described with minor modifications.^46–48^ One hundred and ninety-one 4-month-old male and female WT and App-KI mice were assigned to six groups (male WT vehicle, n = 14; male WT suvorexant, n = 17; male WT lemborexant, n = 15; male App-KI vehicle, n = 19; male App-KI suvorexant, n = 18; male App-KI lemborexant, n = 19; female WT vehicle, n = 15; female WT suvorexant, n = 16; female WT lemborexant, n = 14; female App-KI vehicle, n = 15; female App-KI suvorexant, n = 14; female App-KI lemborexant, n = 15). Mice were orally treated with vehicle, suvorexant (30 mg/kg), or lemborexant (30 mg/kg) once daily for 60 d. In this study, drug-treated 6-month-old WT and App-KI mice were used. Each arm of the Y-maze apparatus was 40 cm long, 12 cm high, 3 cm wide at the bottom, and 10 cm wide at the top. The arms converged to form an equilateral triangular central area measuring 4 cm along the longest axis. Each mouse was individually placed on one of the three arms of the apparatus and allowed to move freely through the maze during an 8-min session under illumination conditions (30−40 lx). The series of arm entries and alteration behavior were counted and analyzed using ANY-maze software (Stoelting, IL, USA). Three batch operations were conducted.

### Immunohistochemistry

Immunohistochemistry was performed as previously described with minor modifications.^49^ After the Y-maze test, six mice per group were used for immunohistochemistry. Mice were anesthetized intraperitoneally with medetomidine, midazolam, and butorphanol and perfused The brains were removed, postfixed in the same fixative and cryoprotected in 20% sucrose, followed by 30% sucrose in 0.1 M phosphate buffer. The 25 μm thick coronal brain slices were cut on a cryostat (CM3050S; Leica Biosystems, Tokyo, Japan). These slices were placed in 0.1 M phosphate buffer, washed three times, fixed in 4% paraformaldehyde in 0.1 M phosphate buffer for 5 min, and permeabilized with 0.1% Triton X-100 in 0.1 M phosphate buffer for 10 min. After incubation in Blocking One Histo (Nacalai, Kyoto, Japan) for 60 min, purified anti-β-amyloid, 1−16 antibody (1:1,000; BioLegend, San Diego, CA, USA) diluted in 0.1 M phosphate buffer was applied to the sections, which were then incubated overnight at 4°C. After three washes in 0.1 M phosphate buffer, goat anti-mouse Alexa Fluor 594 (1:10,000; Thermo Fisher Scientific) and Cellstain^®^-Hoechst 33342 solution (1:2,000; DOJINDO, Kumamoto, Japan) diluted in 0.1 M of phosphate buffer were added to the sections for 1 h at 20−30°C. After three washes in 0.1 M phosphate buffer, the sections were mounted on a glass slide coated with MAS (Matsunami, Osaka, Japan) using Fluorescence Mounting Medium (Dako, Santa Clara, CA, USA) and a coverslip. Fluorescent images were captured using an all-in-one fluorescence microscope (BZ-X800; KEYENCE, Osaka, Japan). The Hip CA1 region was identified using the Mouse Brain Atlas (Franklin and Paxinos, 1997). For quantitative immunohistochemistry analysis, six images were acquired from one sample and analyzed using Hybrid Cell Count software (KEYENCE). After creating regions of interest in the Hip CA1 region, positive regions with signal intensity above the background threshold and >28 µm^2^ were defined as Aβ-positive areas and aggregates to exclude nonspecific signals.

### Statistical analysis

Data are presented as mean + SD for Figures 3 and 4, Supplemental Figures 1 and 2, and mean ± SEM for the others. Statistical analyses were performed using GraphPad Prism 6 software (GraphPad Software, San Diego, CA, USA). Statistical significance was determined using one-way or two-way analysis of variance, and the Tukey−Kramer multiple comparison test was used for post hoc analysis when F ratios were significant. All data and results of the statistical analyses are presented in Supplemental Table 1.

## Results

### Expression of *OXR* gene in the mouse brain

The distribution of the *OXR* gene has been described,^50,51^ but its expression in target week-old mice is unknown. Therefore, we examined the expression of *OX1R* and *OX2R* in the Hip, OFC, cerebral cortex, striatum, nucleus accumbens, and LHA of 6-month-old male WT mice. The results showed that *OX1R* and *OX2R* were expressed in all brain regions (Figure 1A, B). Furthermore, *OX1R* and *OX2R* were most highly expressed in LHA among these brain regions.

**Figure 1.**
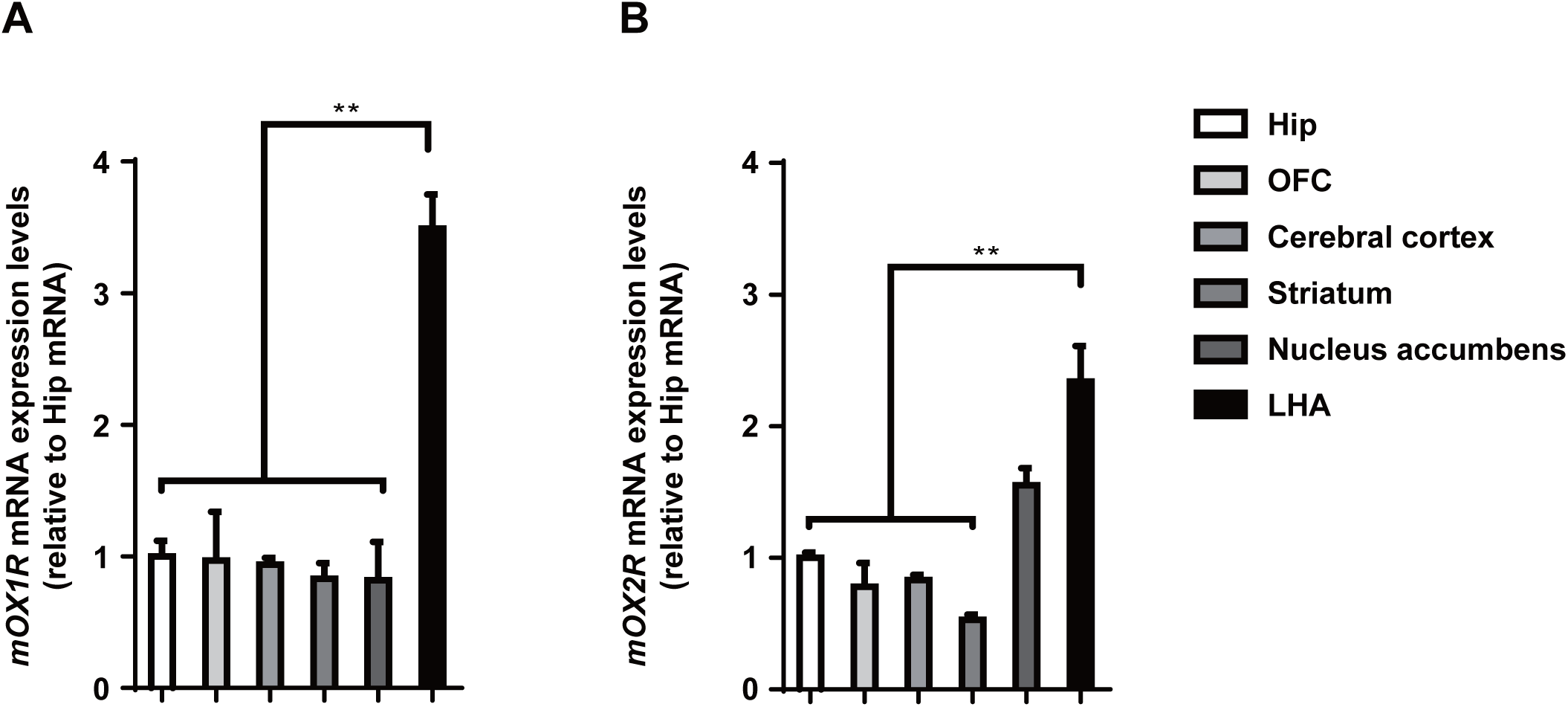
Expression of *OX1R* and *OX2R* gene in the 6-month-old WT male mice. (A) *OX1R* mRNA levels and (B) *OX2R* mRNA levels in the hippocampus (Hip), orbitofrontal cortex (OFC), cerebral cortex, striatum, nucleus accumbens, and lateral hypothalamic area (LHA). ** *p* < 0.01 compared to the respective expression level in Hip. All data were analyzed by one-way ANOVA and expressed as means ± SEM (n = 4 mice).

### Brain levels of OXR antagonists

Understanding the brain pharmacokinetics is important for determining drug dosing schedules and monitoring drug efficacy and side effects.^52^ The pharmacokinetics of suvorexant and lemborexant in plasma, cerebrospinal fluid, and brain have been partially reported.^53–57^ However, reports on the pharmacokinetics of lemborexant in the brain and the same dose of suvorexant or lemborexant in the target week-old mice are lacking. Early Aβ deposition in the Hip and OFC has been reported in patients with MCI.^58^ In addition, early metabolic abnormalities have been reported in the LHA of the AD mouse model.^59,60^ The LHA is also an important site for evaluating the effects of OX receptor antagonists in the early stages of AD mouse models, as it is known to contain OX neuronal cell bodies.^10^ Furthermore, we confirmed the expression of *OX1R* and *OX2R* in the Hip, OFC, and LHA (Figure 1A, B). Therefore, the levels of suvorexant and lemborexant in the brain were examined in 6-month-old male WT mice. Following single oral drug administration, drug levels increased in a time-dependent manner in each brain region, reaching maximum levels (suvorexant: Hip, 8.65 nM at 40 min; OFC, 13.53 nM at 20 min; LHA, 9.94 nM at 40 min and lemborexant: Hip, 30.52 nM at 15 min; OFC, 17.56 nM at 15 min; LHA, 32.07 nM at 15 min) and then decreased (Figure 2A, B). Additionally, brain drug levels 24 h after administration were as follows: Hip, 5.64 nM; OFC, 6.88 nM; LHA, 4.82 nM in suvorexant (Figure 2C) and Hip, 10.56 nM; OFC, 3.01 nM; LHA, 1.64 nM in lemborexant (Figure 2D). The pharmacokinetic parameters of suvorexant and lemborexant in the Hip, OFC, and, LHA are shown in Table 2.

**Figure 2.**
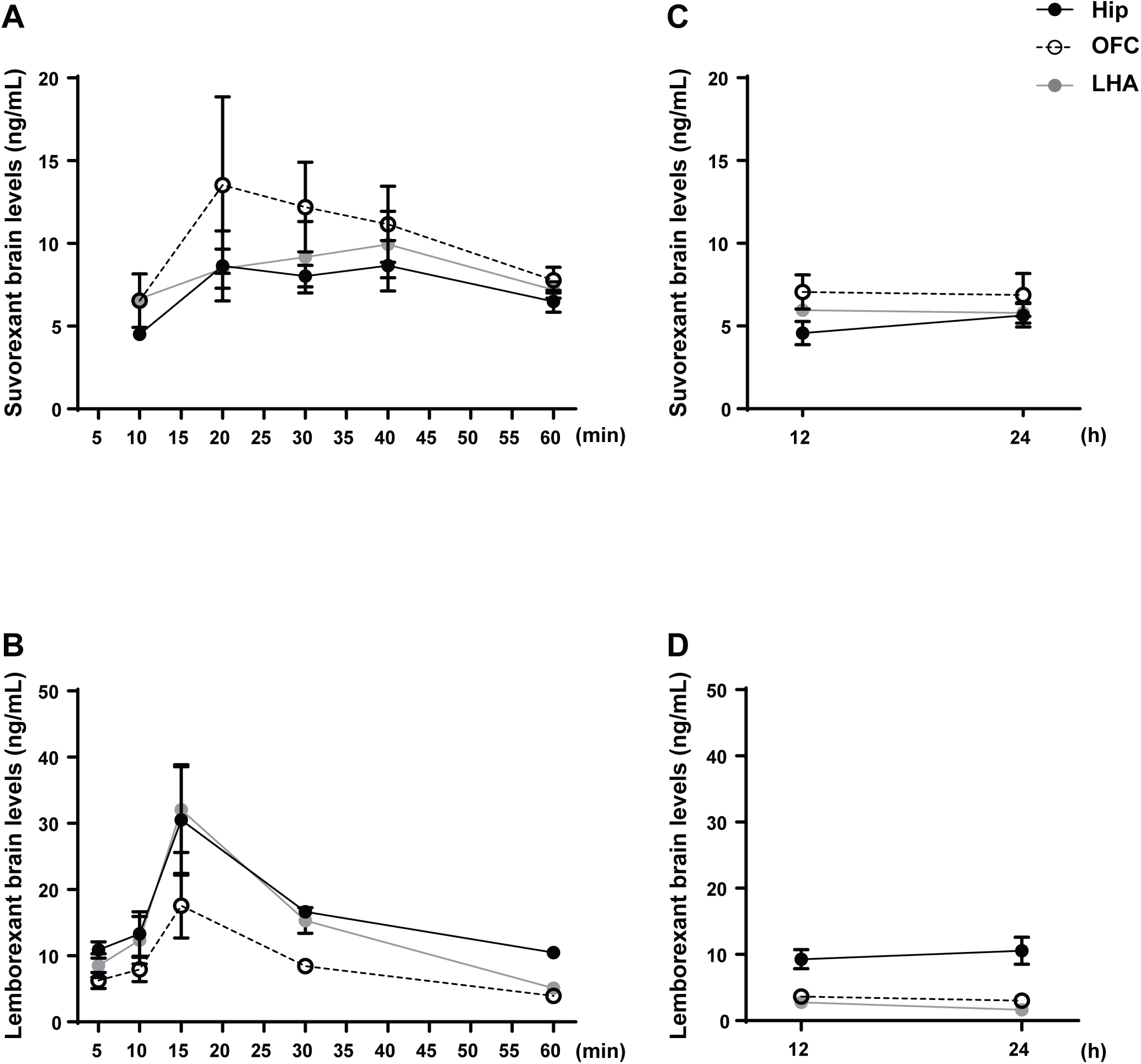
Brain level-time curves for suvorexant and lemborexant in 6-month-old WT male mice. (A) Suvorexant level-time curves at 10−60 min and (B) 12 and 24 h after a single oral dose of suvorexant (30 mg/kg). (C) Lemborexant level-time curves at 5−60 min and (D) 12 and 24 h after a single oral dose of lemborexant (30 mg/kg). All data are expressed as means ± SEM (n = 5 mice at each time point). Hip, hippocampus; OFC, orbitofrontal cortex; LHA, lateral hypothalamic area.

**Table 2.**
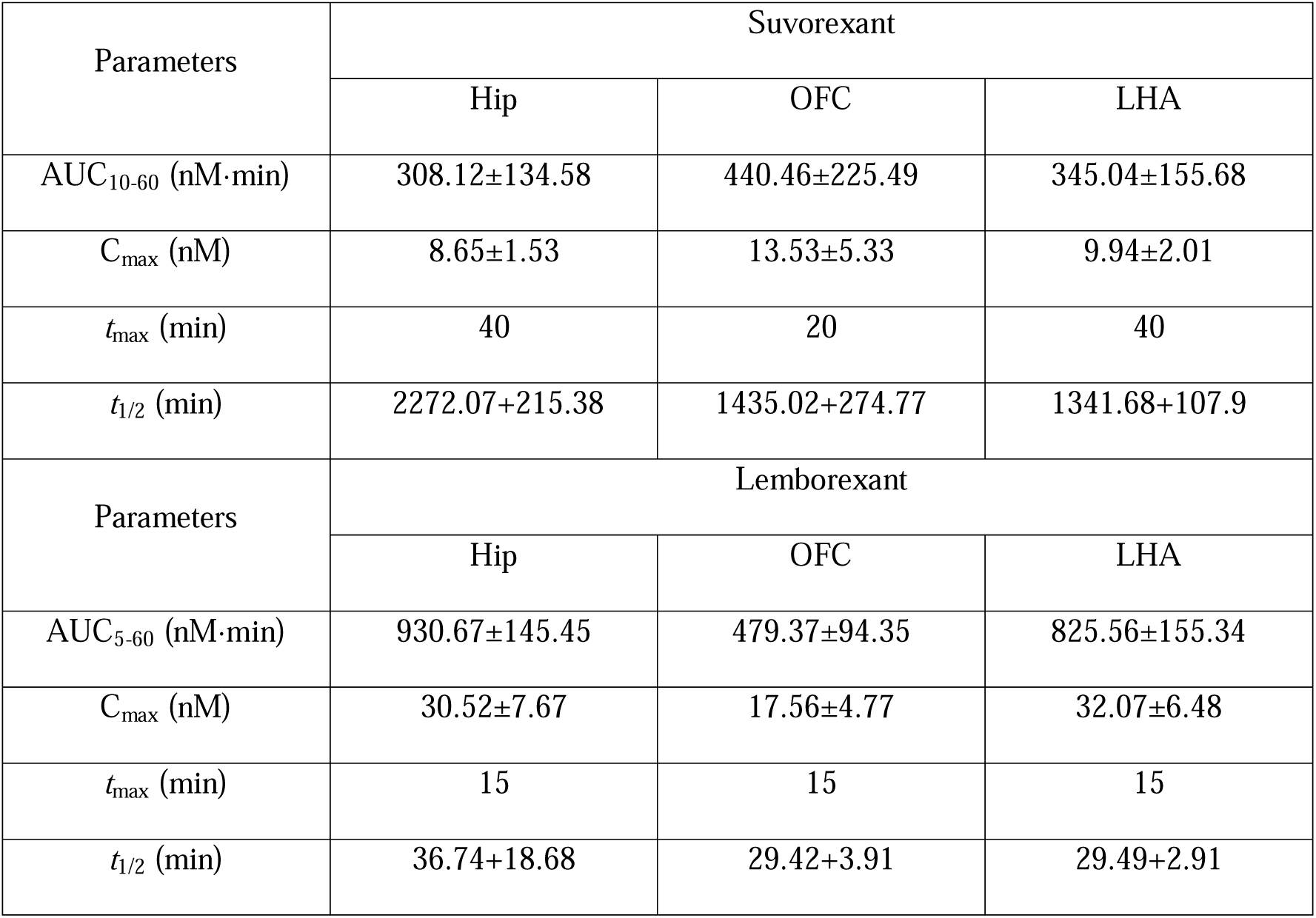
Pharmacokinetic parameters for suvorexant and lemborexant in mice brain.

### Effect of OXR antagonist on spontaneous locomotor activity

In this study, we identified the pharmacokinetics of suvorexant and lemborexant in the Hip, OFC, and LHA (Figure 2). However, assessing the brain pharmacokinetics and their effects and side effects is important. The effects of suvorexant and lemborexant on locomotor activity have already been reported.^29,54^ However, the effects of suvorexant and lemborexant on locomotor activity 24 h after drug administration and in 6-month-old male WT mice have not been reported. Therefore, we administered a single dose of vehicle, suvorexant (30 mg/kg, p.o.), or lemborexant (30 mg/kg, p.o.) to 6-month-old male WT mice and assessed their locomotor activity immediately and 24 h after administration. In the results measured immediately after administration, locomotor activity decreased in a time-dependent manner in all three groups and was significantly faster in the WT suvorexant and WT lemborexant than in the WT vehicle (Figure 3A, B). Furthermore, WT lemborexant showed a significantly faster decrease in locomotor activity than WT suvorexant. In addition, no significant differences were noted in locomotor activity between the three groups 24 h after administration (Figure 3C, D).

**Figure 3.**
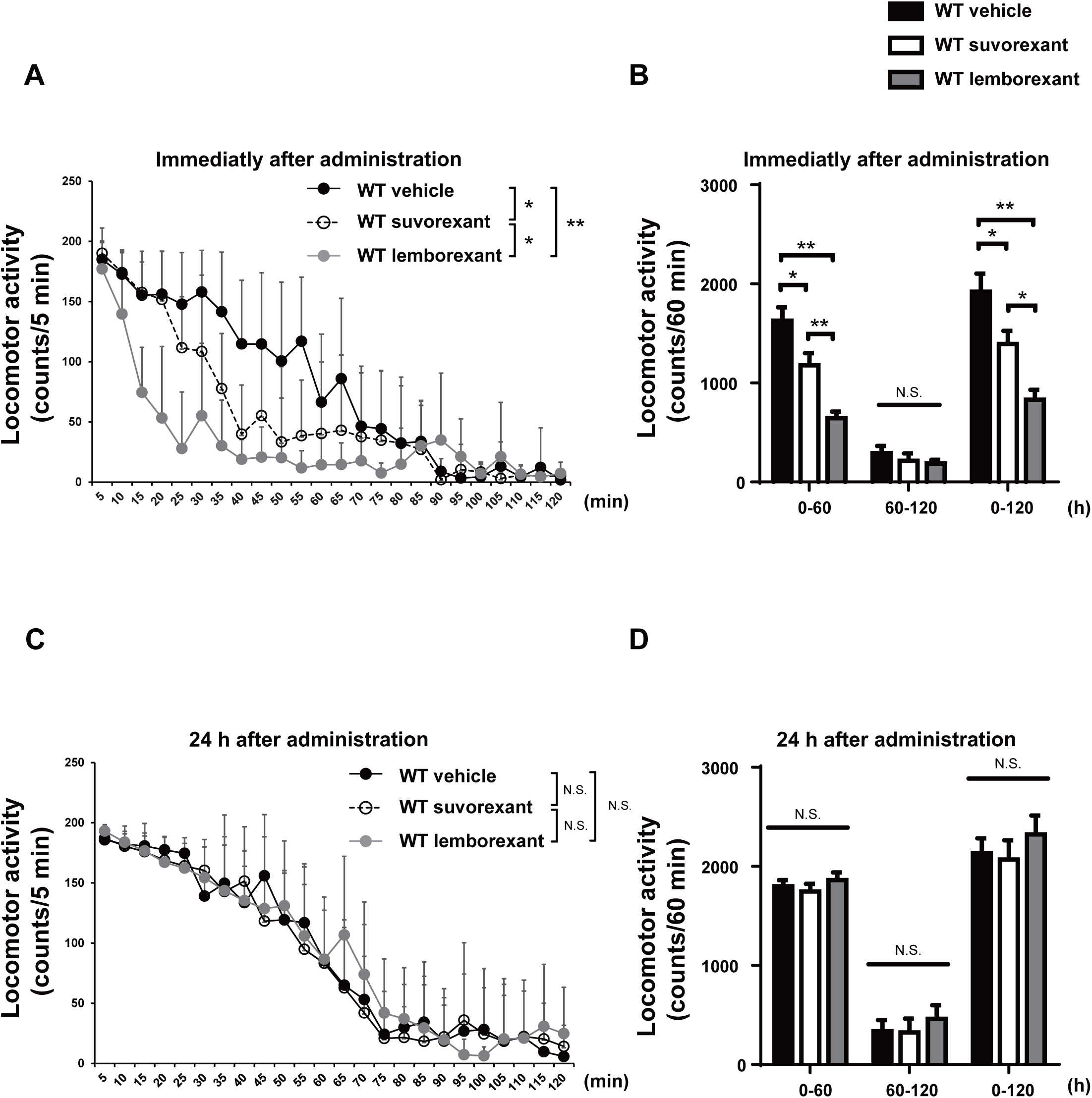
Effect of suvorexant and lemborexant on spontaneous locomotor activity in 6-month-old WT male mice. (A) Time-course changes in locomotor activity from 0 to 120 min after a single oral dose administration of vehicle, suvorexant, or lemborexant. (B) Locomotor activity for 0−60, 60−120 min, and 0−120 min after a single oral dose administration of vehicle, suvorexant, or lemborexant. (C) Time course of locomotor activity from 0 to 120 min 24 h after oral administration of vehicle, suvorexant, or lemborexant. (D) Locomotor activity for 0−60, 60−120 min, and 0−120 min 24 h after a single oral dose administration of vehicle, suvorexant, or lemborexant. * *p* < 0.05, ** *p* < 0.01. All data are expressed as means + SD (n = 9 mice for each group).

### Effect of OXR antagonist in the early stage of AD model mice induced cognitive impairment

Suvorexant and almorexant have been reported to suppress cognitive impairment in mouse models of AD.^32,33^ Therefore, we investigated the effect of suvorexant and lemborexant on the cognitive impairment observed in App-KI mice, a second-generation mouse model of AD that closely mimics AD pathology by harboring Swedish, Iberian, and Arctic mutations in the *amyloid precursor protein* gene, and has been reported to show cognitive impairment.^37,61,62^ The 4-month-old male and female WT and App-KI mice were assigned to six groups (WT vehicle, WT suvorexant, WT lemborexant, App-KI vehicle, App-KI suvorexant, and App-KI lemborexant) and vehicle, suvorexant (30 mg/kg), or lemborexant (30 mg/kg) was administered orally once a day for 60 d. No group showed any differences in changes in body weight during administration (Figure 4A). After 60 d, cognition was assessed using the Y-maze test. During the 8-min session, the WT vehicle showed approximately 14 arm entries and 72% spontaneous alternation, while the App-KI vehicle showed approximately 12 arm entries; however, spontaneous alternation was significantly lower at 55.8% (Figure 4B, C). For WT suvorexant and WT lemborexant, the arm entries were approximately 16 and 15, respectively, and spontaneous alternation rates were 68.6% and 69.4%, respectively, with no significant differences between them and the WT vehicle. App-KI suvorexant and App-KI lemborexant had approximately 13 arm entries each, which was not significantly differences from the App-KI vehicle; however, spontaneous alternation was significantly higher than that of the App-KI vehicle at 68.8% and 68.6%, respectively. Similar results were obtained when sex differences were examined (Supplementary Figures 1 and 2).

**Figure 4.**
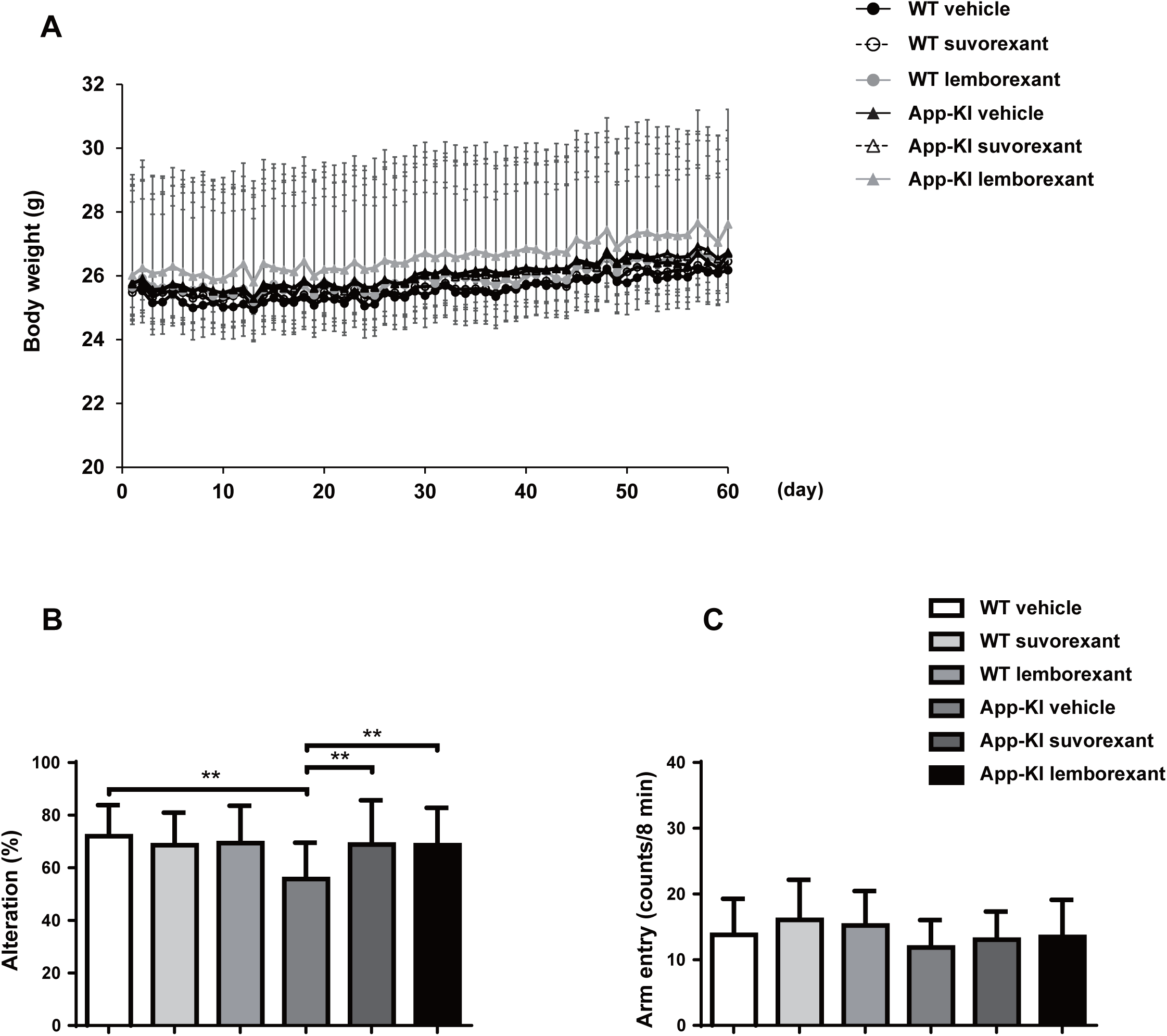
Effect of suvorexant and lemborexant on cognitive impairment in 6-month-old WT and App-KI mice. (A) Sequential changes in body weight during the administration period of vehicle, suvorexant, or lemborexant for 1–60 d. Y-maze test (B). Alteration behavior (C). Total arm entries. ** *p* < 0.01. All data are expressed as means + SD (n = 29 for WT vehicle, n = 33 for WT suvorexant, n = 29 for WT lemborexant, n = 34 for App-KI vehicle, n = 32 for App-KI suvorexant, n = 34 for App-KI lemborexant).

### Effect of OXR antagonist on A**β** accumulation

Aβ accumulation has been reported to correlate with the progression of AD and MCI pathologies and is used for diagnosis.^63^ Suvorexant and almorexant have been reported to inhibit Aβ accumulation in the brains of AD mouse models.^30,32^ App-KI mice develop Aβ accumulation in the CA1 region of the Hip at 3–4 months of age.^64^ Therefore, the assessment of Aβ accumulation in the Hip CA1 region is useful for the pathological evaluation of AD. Aβ accumulation in the Hip CA1 region was assessed in mice after behavioral analysis, and the staining area and number of Aβ aggregates were measured. The WT vehicle, WT suvorexant, and WT lemborexant showed no Aβ accumulation (Figure 5A). The App-KI vehicle showed 4.18% Aβ positive areas and about 58 number of Aβ aggregates. Meanwhile, App-KI suvorexant and App-KI lemborexant showed 1.42% and 0.98% Aβ positive areas, respectively and approximately 34 and 28 the number of Aβ aggregates, respectively, and compared to App-KI vehicle showed Aβ positive areas and the number of Aβ aggregates were found to be significantly decreased (Figure 5B, C).

**Figure 5.**
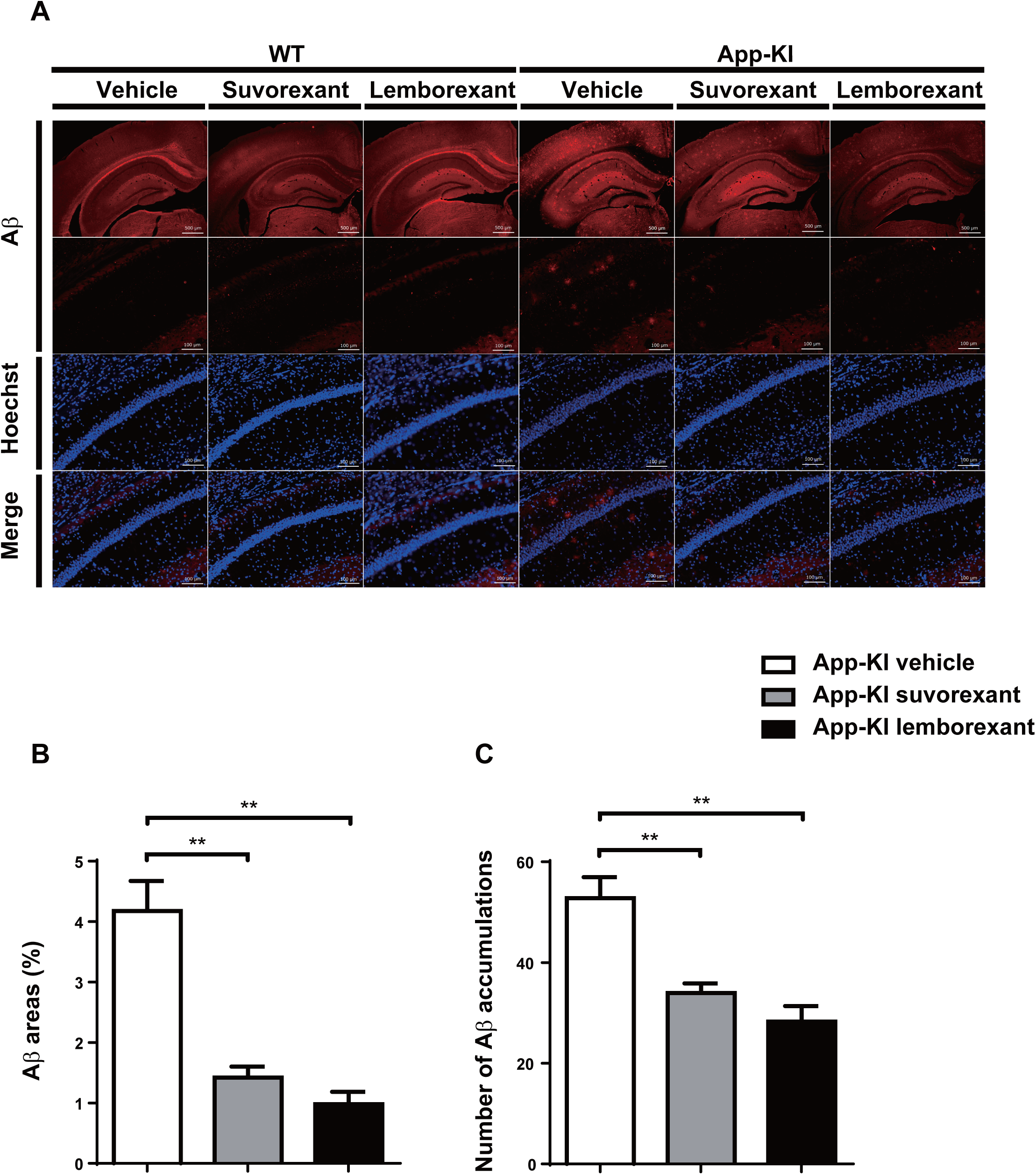
Effect of suvorexant and lemborexant on Aβ accumulation in 6-month-old App-KI mice. Immunohistochemistry. (A) Representative images of the hippocampal CA1 region. Measurement (B) area and (C) number of Aβ debris. ** *p* < 0.01. All data are expressed as means ± SEM (n = 6 mice for each group, 6-images/sample). Scale bar = 500 μm and 100 μm.

## Discussion

In this experiment, we showed that *OX1R* and *OX2R* were expressed in the Hip, OFC, cerebral cortex, striatum, nucleus accumbens, and LHA tissues. These results are consistent with previously reported *in situ* hybridization results for the *OX1R* and *OX2R* genes.^50,51,65^ These brain regions are innervated by OX nerves,^12,66^ and have been implicated in clock and circadian oscillations, memory, decision making, feeding, and motor activity.^67–73^ Furthermore, the *OX1R* and *OX2R* genes were most highly expressed in the LHA between these brain regions in 6-month-old male WT mice. In addition, endogenous OX release at the LHA has an excitatory effect and may lead to the amplification of excitation by further activating other OX neurons.^74^ These considerations made it reasonable to assess the efficacy of OXR antagonists in these brain regions in 6-month-old male WT mice.

Suvorexant and lemborexant were administered as single oral doses to 6-month-old male WT mice, and their pharmacokinetics in the Hip, OFC, and LHA were investigated. The results of the brain pharmacokinetic analysis of suvorexant and lemborexant showed slight differences between tissues, but the highest levels were detected after 20−40 min for suvorexant (after 20 min for OFC, after 40 min for Hip and LHA) and 15 min for lemborexant in all three brain regions. These results were not entirely consistent with previously reported plasma drug level-time profiles.^57,75^ This may be due to differences in postabsorptive drug transfer and tissue excretion. Furthermore, in suvorexant, drug levels above Ki values of OX1R and OX2R (OX1R, 0.55 nM; OX2R, 0.35 nM) were detected in the Hip, OFC, and LHA up to 24 h after administration. Meanwhile, drug levels above the Ki values of OX1R and OX2R (OX1R, 8.1 nM; OX2R, 0.48 nM) were detected only in the Hip at 24 h after administration in lemborexant, but only OX2R was detected above Ki values in the OFC and LHA. These results suggest that suvorexant or lemborexant administered orally at 30 mg/kg may be effective in the brain 24 h after administration.

Regarding the effects of vehicle, suvorexant, or lemborexant on locomotor activity in 6-month-old male WT mice, all three groups showed a chronological decrease in locomotor activity immediately after administration; however, WT suvorexant and WT lemborexant showed a significant decrease compared to the WT vehicle. Furthermore, WT lemborexant caused a significantly faster decrease in locomotor activity than WT suvorexant. This suggests that lemborexant may be more effective in inducing sleep than suvorexant at the same dose. Previous reports have shown that lemborexant is more effective than suvorexant in improving sleep onset than suvorexant.^76^ However, no effect was observed in locomotor activity after 24 h. No significant differences were noted between the control and OXR antagonist-treated groups in awake time, non-REM sleep time, and REM sleep time 24 h after suvorexant administration and 12 h after lemborexant administration.^77,78^ Combined with the results of the brain pharmacokinetic analysis, the drug was still effective in the brain after 24 h administration but did not affect locomotor activity. Therefore, administering 30 mg/kg orally once a day to assess its effects on cognitive function seems reasonable. However, AD is a neurodegenerative disease; therefore, repeated drug administration has been used to assess its effects on cognitive function. In the Y-maze study, cognitive function was significantly impaired in the App-KI vehicle compared to the WT vehicle, consistent with previous reports.^37,61^ Furthermore, repeated administration of suvorexant or lemborexant significantly improved cognitive impairment observed in the App-KI vehicle. These results are consistent with those of previous reports of improved cognitive impairment in AD models after repeated administration of suvorexant or almorexant.^32,33^ Furthermore, suvorexant or lemborexant had no effect on arm entry. The results for arm entry in the Y-maze test and locomotor activity showed that locomotor activity was not only observed after 24 h of a single administration, but also did not have an effect on locomotor activity after 24 h of repeated administration. These results suggest that suvorexant and lemborexant inhibit the pathophysiology of AD without affecting motor activity 24 h after administration and ameliorate cognitive impairment in the early stages of AD. In this study, the effects of suvorexant and lemborexant were only examined using the Y-maze test, but App-KI mice have also been reported to show impairments in other tests that assess cognitive function, such as the novel object recognition test, water maze test, and contextual fear conditioning.^79^ Therefore, the effects of suvorexant and lemborexant should be examined in other behavioral analyses.

Immunohistochemistry revealed that the suvorexant or lemborexant repeated administration improved the accumulation of Aβ in the Hip CA1 region. Suvorexant and lemborexant induce sleep by antagonizing the G protein-coupled receptors (OX1R and OX2R). Therefore, the decrease in Aβ in the Hip CA1 region may also be mainly due to OX1R and OX2R antagonism. An et al. reported that OX inhibits phagocytosis and degradation of Aβ fibrils by microglial cells by decreasing molecules, such as PI3K, Akt, and p38-MAPK, that regulate phagocytosis. ^80^ Therefore, suvorexant and lemborexant might activate microglial phagocytosis. Furthermore, during sleep, debris, such as Aβ, tau protein, and misfolded proteins, in the brain are removed by the glymphatic pathway.^16,23,25,81^ Therefore, the glymphatic pathway is believed to be activated by the repeated administration of suvorexant or lemborexant.

In conclusion, the results of this study revealed the expression of the *OX1R* and *OX2R* genes in the brain, suvorexant and lemborexant on brain pharmacokinetics and the effect on locomotor activity in 6-month-old male WT mice. Therefore, the efficacy of a single oral dose of 30 mg/kg and its safety after 24 h were assessed, providing new information on the use of suvorexant and lemborexant. Additionally, our results showed that repeated administration of suvorexant or lemborexant improved cognitive impairment in App-KI mice. Abnormalities in neurotransmitters (dopamine, serotonin, glutamate, GABA, histamine, adrenaline, noradrenaline, glycine, and acetylcholine) are some of the causes of cognitive impairment,^82–86^ and OXR antagonists may have improved them. OX has been reported to regulate dopamine, serotonin, GABA, and glutamate neurons, among others^87–89^; these nerves have also been discussed for their involvement in AD pathology,^90–93^ and suvorexant and lemborexant may indirectly normalize their neural functions. Therefore, whether these symptoms improve with repeated administration of suvorexant or lemborexant must be investigated. In addition, repeated administration of suvorexant or lemborexant also suppressed Aβ deposition in the Hip CA1 region of early-stage AD model mice, which may also ameliorate previously reported synaptic morphological and electrophysiological changes in the Hip CA1 region, the turnover of proteins associated with presynaptic terminals previously reported in App-KI mice.^64,94–96^ In the study did not find any significant differences between suvorexant and lemborexant, except for locomotor activity outcomes. This suggests that these differences cannot be assessed by cognitive function assessment or Aβ quantification and that electrophysiology and sleep-related factors need to be evaluated. Furthermore, this study showed that suvorexant and lemborexant effectively suppressed the pathology of MCI and early AD. However, their effects on the intermediate and late stages of AD remain unclear and require further investigation. Based on these considerations, further studies on factors associated with cognitive dysfunction and Aβ reduction in this study will elucidate the regulatory mechanisms of AD pathology through OX neurons and lead to new therapeutic approaches for patients with early-stage AD.

## Acknowledgments

We would like to thank Editage (www.editage.jp) for English language editing.

## Funding

This study was supported by Grant of AMED, Japan (JP21wm0425014); The Clinical Research Promotion Foundation (2022); Ichihara International Scholarship Foundation (2022); Institute of Pharmaceutical Life Sciences, Aichi Gakuin University (2022-23); THE HORI SCIENCE AND ARTS FOUNDATION (2023), The Nitto Foundation (2024), THE FORDAYS FOUNDATION (2024) Grant-in-Aid for Early-Career Scientists (23K14364; 2023-2026).

## Author Contributions

**Kazuhiro Hada** (Conceptualization; Data curation; Formal analysis; Funding acquisition; Investigation; Methodology; Project administarion; Resources; Supervision; Visualizatio; Writing-original draft); **Yuki Murata** (Methodology; Resources; Writing-review and editing); **Ohi Yoshiaki** (Funding acquisition; Supervision; Writing-review and editing); **Sana Hashimoto** (Investigation); **Hinata Watanabe** (Investigation); **Kayoko Ozeki** (Supervision; Writing-review and editing); **Takaomi C Saido** (Resources; Writing-review and editing); **Takashi Saito;** (Resources; Writing-review and editing); **Hiroki Sasaguri** (Resources; Writing-review and editing); **Hiroyuki Mizoguchi** (Funding acquisition; Supervision; Writing-review and editing); **Kiyofumi Yamada** (Funding acquisition; Supervision; Writing-review and editing) **Yoshifumi Wakiya** (Supervision; Writing-review and editing)

## Declaration of conflicting interests

Takaomi C Saido is an Editorial Board Member of this journal but was not involved in the peer-review process of this article nor had access to any information regarding its peer-review.

## Data availability statement

The data supporting the findings of this study are available within the article and/or its supplemental material.

---

**Supplemental Figure 1.**
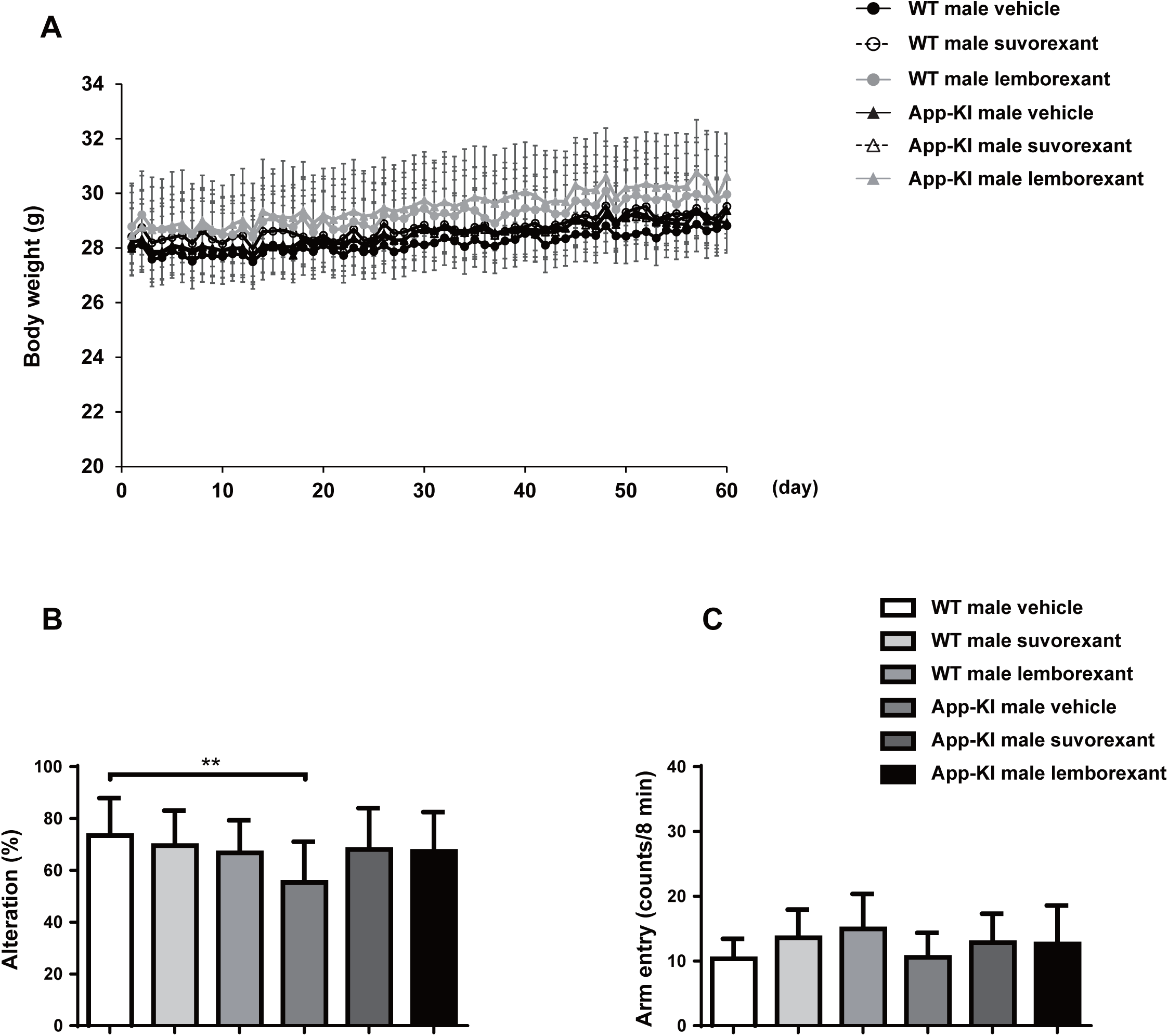
Effect of suvorexant and lemborexant on spontaneous locomotor activity in 6-month-old WT and App-KI male mice. (A) Sequential changes in body weight during the administration period of vehicle, suvorexant, or lemborexant for 1–60 d. Y-maze test (B). Alteration behavior (C). Total arm entry. * *p* < 0.01. All data are expressed as means + SD (n = 14 for WT vehicle, n = 17 for WT suvorexant, n = 15 for WT lemborexant, n = 19 for App-KI vehicle, n = 18 for App-KI suvorexant, n = 19 for App-KI lemborexant).

**Supplemental Figure 2.**
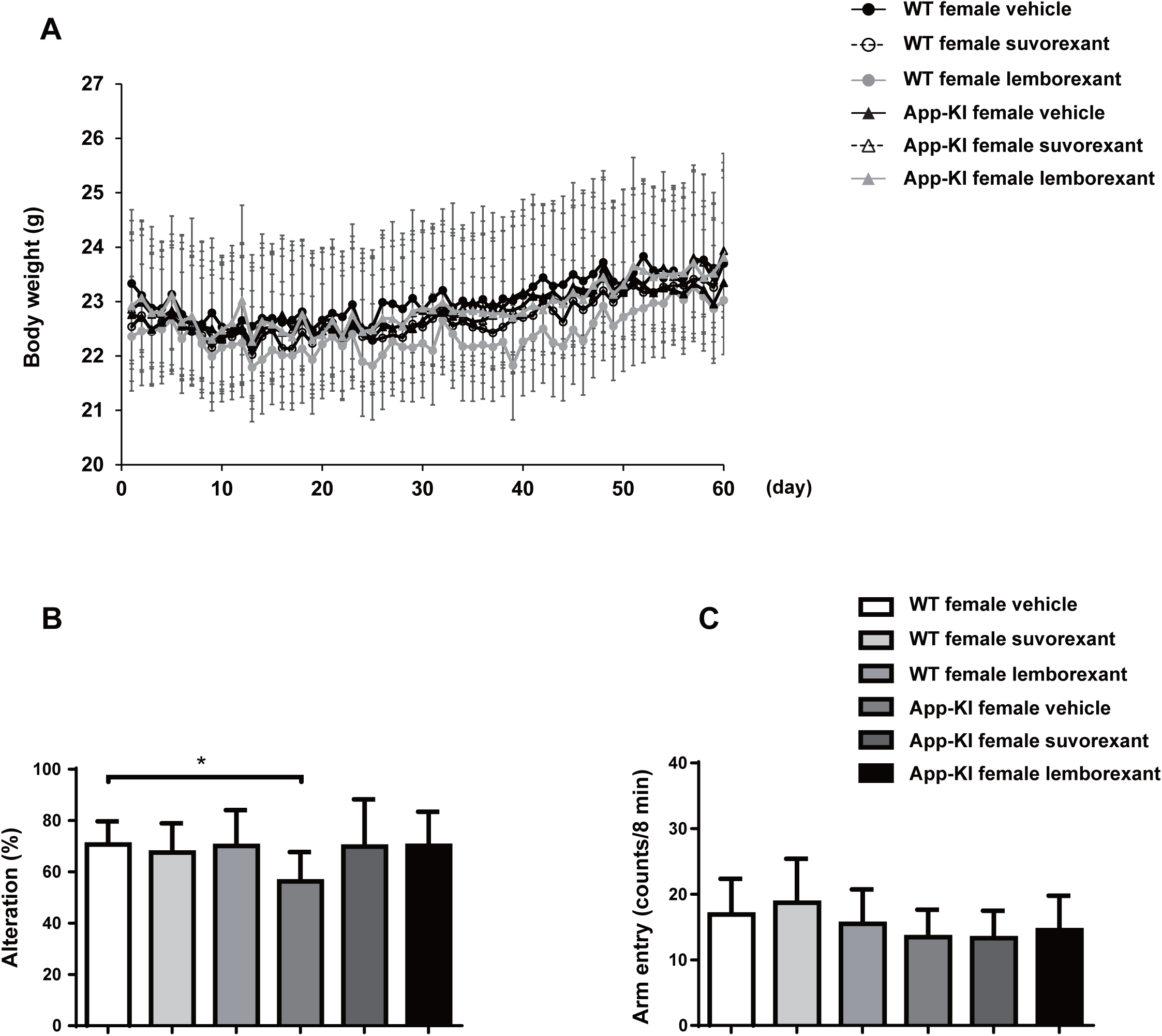
Effect of suvorexant and lemborexant on spontaneous locomotor activity in 6-month-old WT and App-KI female mice. (A) Sequential body weight changes during the administration period of vehicle, suvorexant, or lemborexant for 1–60 d. Y-maze test (B). Alteration behavior (C). Total arm entry. * *p* < 0.05. All data are expressed as means + SD (n = 15 for WT vehicle, n = 16 for WT suvorexant, n = 14 for WT lemborexant, n = 15 for App-KI vehicle, n = 14 for App-KI suvorexant, n = 15 for App-KI lemborexant).

## Notes

### Competing Interest Statement

The authors have declared no competing interest.

